## Supplemental information for "Mapping of morpho-electric features to molecular identity of cortical inhibitory neurons"

### Supporting information

#### S1 Appendix: electrophysiological features list:

AHP_depth

AP_amplitude

AP1_amp

AP_duration_half_width

fast_AHP

voltage_base

AHP_time_from_peak

doublet_ISI

inv_fifth_ISI

inv_first_ISI

inv_fourth_ISI

inv_second_ISI

inv_third_ISI

mean_frequency

**S2 Appendix: NeuroM features list:**

number_of_bifurcations

number_of_forking_points

number_of_sections

number_of_segments

partition

partition_asymmetry

partition_pairs

remote_bifurcation_angles

section_bif_branch_orders

section_bif_radial_distances

section_branch_orders

section_end_distances

section_lengths

section_path_distances

section_radial_distances

section_strahler_orders

section_term_branch_orders

segment_lengths

segment_midpoints

segment_radial_distances

terminal_path_lengths_per_neurite

total_length

**S3 Appendix: common me-features list:**

'ephys_AHP_time_from_peak',

'ephys_AP1_amp',

'ephys_AP_duration_half_width',

'ephys_doublet_ISI',

'ephys_fast_AHP',

'ephys_inv_fifth_ISI',

'ephys_inv_first_ISI',

'ephys_inv_fourth_ISI',

'ephys_inv_second_ISI',

'ephys_inv_third_ISI',

'ephys_mean_frequency',

'ephys_voltage_base',

'morpho_nm_number_of_bifurcations',

'morpho_nm_number_of_forking_points',

'morpho_nm_number_of_sections',

'morpho_nm_number_of_segments',

'morpho_nm_partition',

'morpho_nm_partition_asymmetry',

'morpho_nm_partition_pairs',

'morpho_nm_remote_bifurcation_angles',

'morpho_nm_section_bif_branch_orders',

'morpho_nm_section_bif_radial_distances',

'morpho_nm_section_branch_orders',

'morpho_nm_section_end_distances',

'morpho_nm_section_lengths',

'morpho_nm_section_path_distances',

'morpho_nm_section_radial_distances',

'morpho_nm_section_strahler_orders',

'morpho_nm_section_term_branch_orders',

'morpho_nm_segment_lengths',

'morpho_nm_segment_midpoints',

'morpho_nm_segment_radial_distances',

'morpho_nm_terminal_path_lengths_per_neurite',

'morpho_nm_total_length'

+ 50 morphological moments

**S4 Appendix: Neurite density moments extraction:**

Moments were computed using the same definition as in Snider et al. (2010):

$$m_{j,k} = \sum_{i=1}^{N} x_i^j y_i^k w_i$$

$(x_i, y_i)$ are the 2-D coordinates of the center of the ith segment and $w_i$ is the length of the arbor segment. Hence, the total length of the neurite shaft is given by the 0th order moment:

$$m_0 = m_{0,0} = \sum_{i=1}^{N} w_i.$$

We modified the first equation to define a density moment within a layer $l$ as:

$$m^{(l)}_{j,k} = \sum_{i=1}^{N} x_i^j y_i^k w_i \quad with\, lim^{(l-1)} < y_i < lim^{(l)}$$

We only considered moments for the special cases of j = 0, k > 0; and of j > 0, k =0 (separating x and y axis)

$$m_k^{x,(l)} = m_{k,0}^{(l)} / \sqrt{m_0} \text{ and}$$

$$m_k^{y,(l)} = m_{0,k}^{(l)} / \sqrt{m_0}$$

with division by $\sqrt{m_0}$ a necessary normalisation in these cases (Snider et al., 2010). We extracted $m_0$, $m_k^{x,(l)}$, $m_k^{y,(l)}$ for $k \in [0, 2]$ and $l \in [1, 5]$ for both BBP and AIBS morphologies. As stated above, $m_0$ gives the total length of the neurite shaft whereas$m_1^{x,y}$ and $m_2^{x,y}$ relate to the mean and standard deviation of the neurite shaft density, respectively, along the $x$ or $y$ axis. Thus, for a given morphology we extracted 5 moments ($m_0^{(l)}$, $m_1^{x,(l)}$, $m_2^{x,(l)}$, $m_1^{y,(l)}$, $m_1^{y,(l)}$) multiplied by 5 layers (*L*1, *L*2/3, *L*4, *L*5, *L*6) multiplied by 2 neurites (*axon, dendrite*) resulting in 50 moments. If a neurite shaft did not extend to a given layer, all moments values were set at zero for that layer.

**S5 Appendix: Brief description of maker densities extraction:**

Briefly, marker expressions from *in-situ* hybridisation experiments were aligned on a reference digital brain and converted to densities using reference values from literature. The reference brain was voxelized and densities of PV, SST and VIP expressing cells were extracted for each voxel across the whole brain. A density for inhibitory cells expressing none of the previously mentioned markers (REST) was also given by the Blue Brain cell atlas.

**S6 Appendix : Formal definition of R-value.**

We implemented the R value as described in Borsos et al., using the same notation:

- $n$ is the number of classes
- $C_i$ the set of instances belonging to class $i$
- $U$the set of all instances $U = C_1 \cup C_2 \cup ... \cup C_n$
- $P_{i,m}$the m-th instance of class $i$
- $\lambda(x) = \{ \begin{smallmatrix} 1, if\ x>0 \\ 0, otherwise \end{smallmatrix}$
- $kNN(P, S)$the subset of $k$-nearest neighbors of instance $P$ that belongs to the set of instance $S$
- θ threshold value above which the instance is considered as belonging to an overlapping region

The *R-value* for a class $i$ is defined as :

$$R(C_i) = \frac{1}{|C_i|} \sum_{m=1}^{|C_i|} \lambda\left(\left|kNN(P_{i,m}, U - C_i\right| - \theta\right)$$

Then, the *R-value* for a dataset f is defined as :

$$R(f) = \frac{1}{|U|} \sum_{i=1}^{n} \sum_{m=1}^{|C_i|} \lambda\left(\left|kNN(P_{i,m}, U - C_i\right| - \theta\right)$$

In their paper, Borsos et al. adapted the *R-value* using two classes $C_{pos}$ and $C_{neg}$, such as $U = C_{pos} \cup C_{neg}$. They introduced the imbalance ratio $IR = \frac{|C_{neg}|}{|C_{pos}|}$and proposed the *augmented R-value*:

$$R(f) = \frac{1}{IR+1}\left(IR \cdot R(C_{neg}) + R(C_{pos})\right)$$

We wanted to be able to use the metric in a case where $n > 2$ (i.e. the dataset is not divided in only $C_{pos}$ and $C_{neg}$). We thus modified the definition of the R value as follow:

$R(f) = \frac{1}{|U|}\sum_{i=1}^{n}\sum_{m=1}^{|C_i|}\lambda\left(\left|kNN(P_{i,m}, U - C_i\right| - \theta_i\right)$, where we fixed the k value as $k = int(0.1\cdot|U|)$ and set $\theta_i = int\left(k/2 \cdot \frac{|U - C_i|}{|U|}\right)$. Thus, the threshold value is corrected to take into account imbalance between classes and can be used for more than 2 classes.

**S1 Table:** Correspondences between m-types or me-types as defined within dataset with common morphological labels

| Common morphological labels | BBP m-types | AIBS me-types |
|---|---|---|
| NGC | NGC | ME_Inh_17, ME_Inh_18, ME_Inh_19, ME_Inh_20 |
| BC | SBC, NBC, LBC | ME_Inh_6, ME_Inh_7, ME_Inh_8, ME_Inh_9, ME_Inh_10, ME_Inh_11, ME_Inh_12, ME_Inh_13, ME_Inh_14, ME_Inh_16 |
| ChC | ChC | ME_Inh_21 |
| MC | MC | ME_Inh_15, ME_Inh_24, ME_Inh_25, ME_Inh_26 |
| BTC | BTC, DBC, BP | ME_Inh_1, ME_Inh_2, ME_Inh_3, ME_Inh_4, ME_Inh_5, ME_Inh_22, ME_Inh_23, ME_Inh_26 |

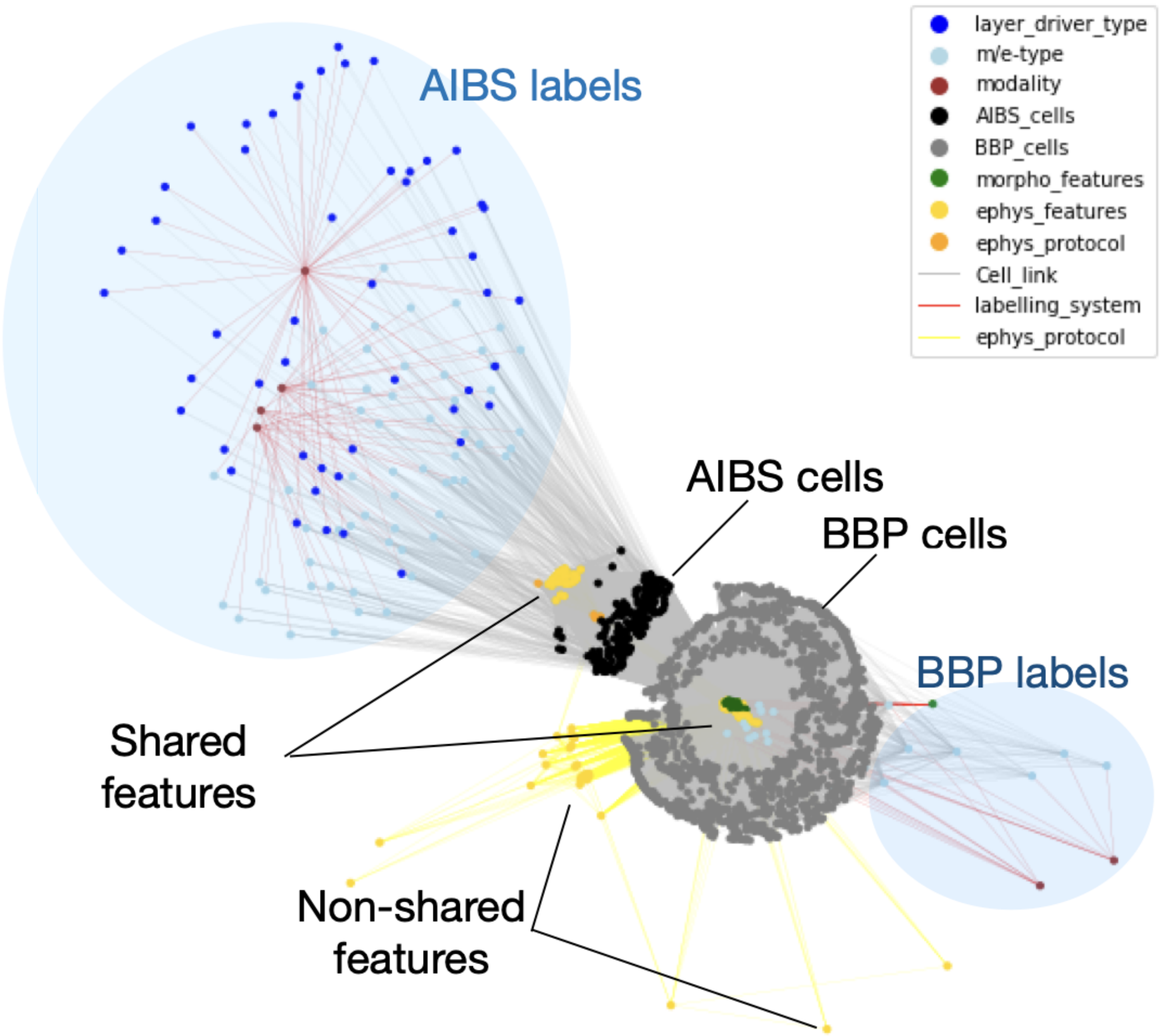


**S1 Figure: Cell-centered "Knowledge Graph"**

We visualized all the pieces of information we had about each cell in a graph. Labels (dark and light blue), modality (e.g. morphological, electrical or genetic classification, dark red), morphological features (green), electrophysiological features (yellow), electrophysiological protocols (gold) were linked to individual cells from either Allen Institute for Brain Science (AIBS) or Blue Brain Project (dataset) for which this information was available.

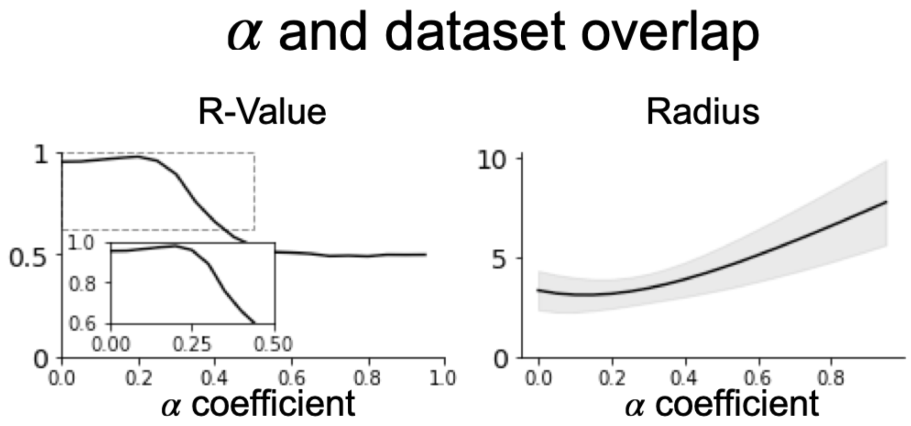


**S2 Figure:** Effect of alpha on datasets overlap (R-value). Left - R-value computed for different values of α. Inset is a zoom of the dashed box. Right - Radius of the global dataset (AIBS +BBP) computed for different values of α. Radius computed as the mean of euclidean distance between each of the global dataset instances. Standard deviation from the euclidean distance between each of the instances are displayed in grey.

### Densities [mm$^{-3}$] along cortical depth

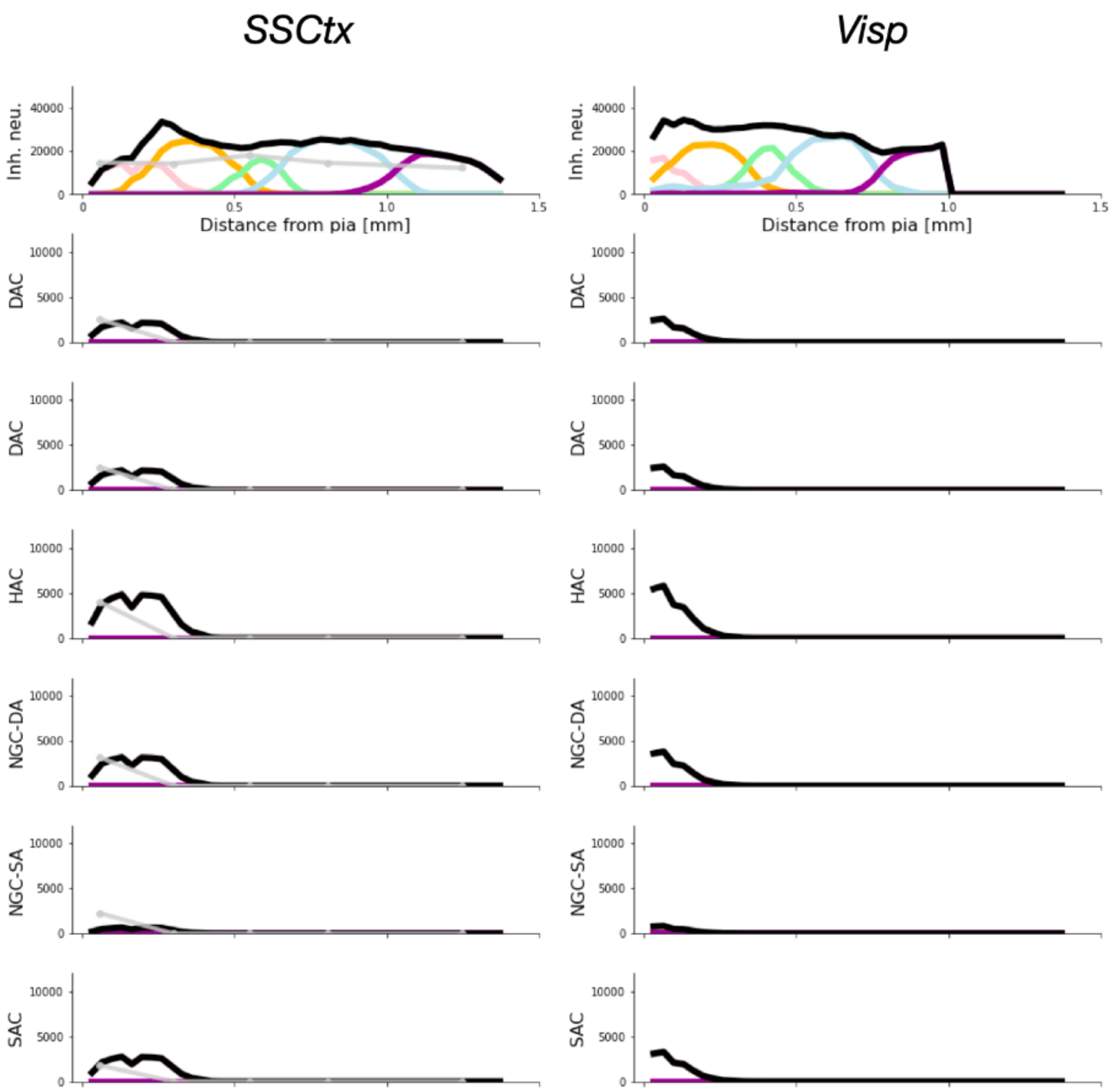

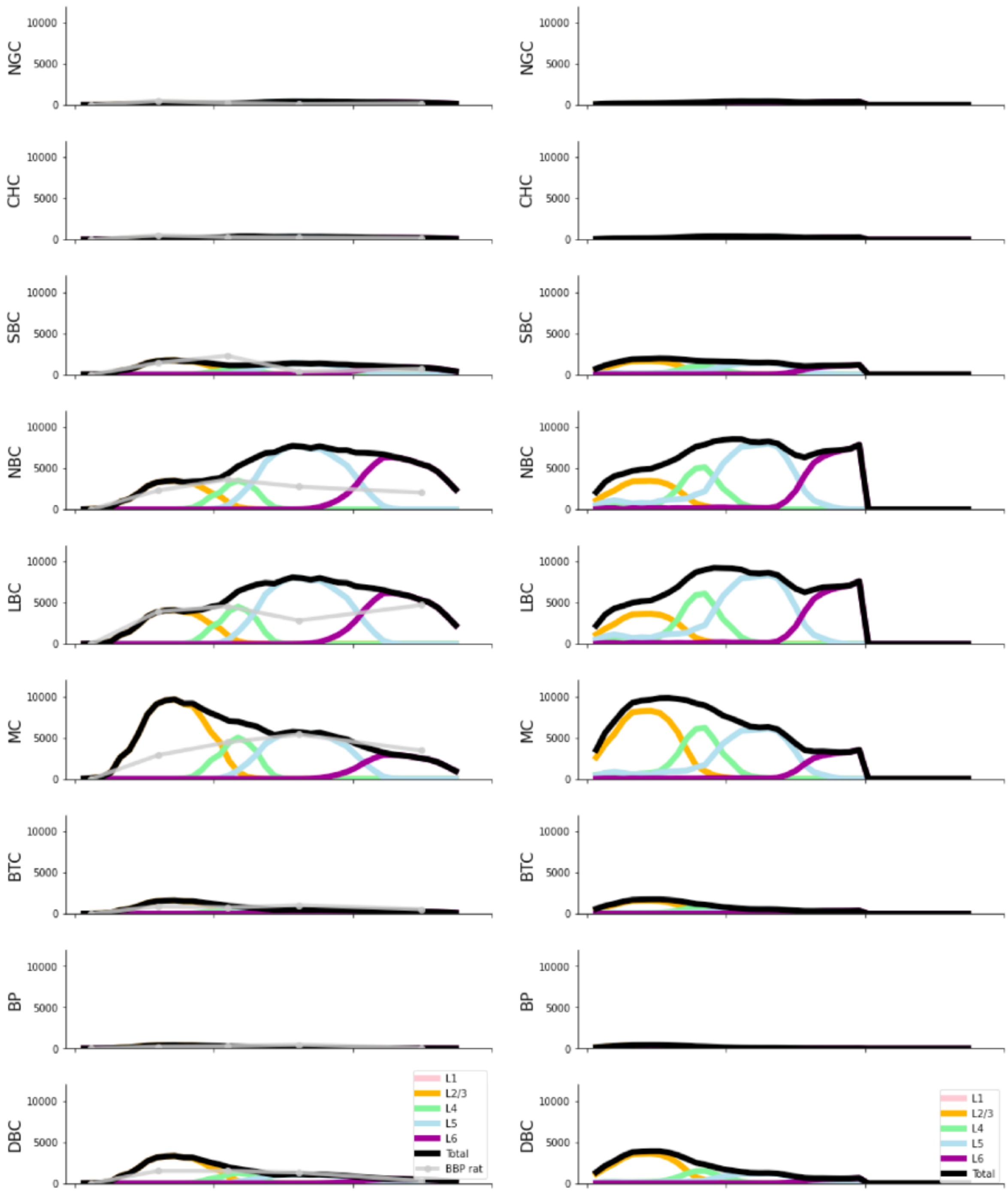


**S3 Figure:** Density profiles of BBP m-types computed from Blue Brain Mouse cell atlas densities combined with probabilistic mapping. Densities are computed as in Fig.5. Left - Profiles for somatosensory cortex (SSCtx). Right - Profiles for visual primary area (Visp). NGC: Neurogliaform cells; CHC: Chandelier cells; SBC: Small Basket cells; NBC: Nested Basket cells; LBC: Large Basket cells; MC: Martinotti cells; BTC: Bitufted cells; BP: Bipolar cells; DBC: Double Bouquet cells.

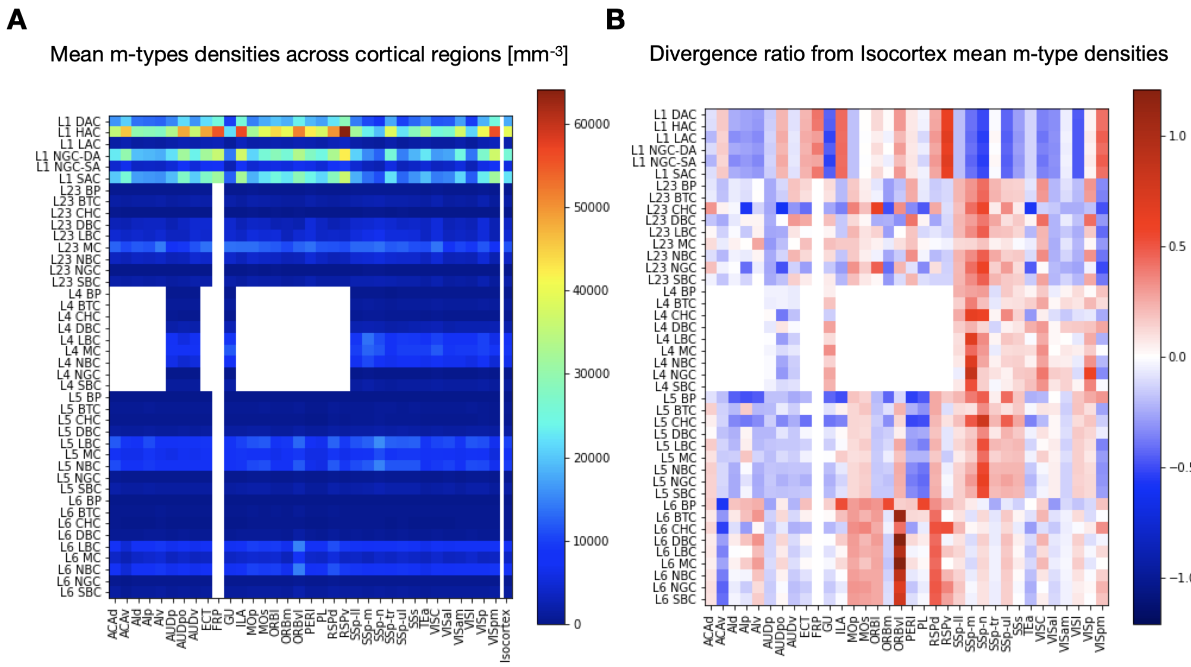


**S4 Figure: A**. Mean m-type densities across cortical regions expressed in $mm^{-3}$. The right column shows mean densities for the whole cortex. **B**. Divergence ratio from the mean density over the whole cortex for each m-type. Ratio was computed as $r^{region}_{m-type} = \frac{mean\ density^{region}_{m-type} - mean\ density^{Isocortex}_{m-type}}{mean\ density^{Isocortex}_{m-type}}$.

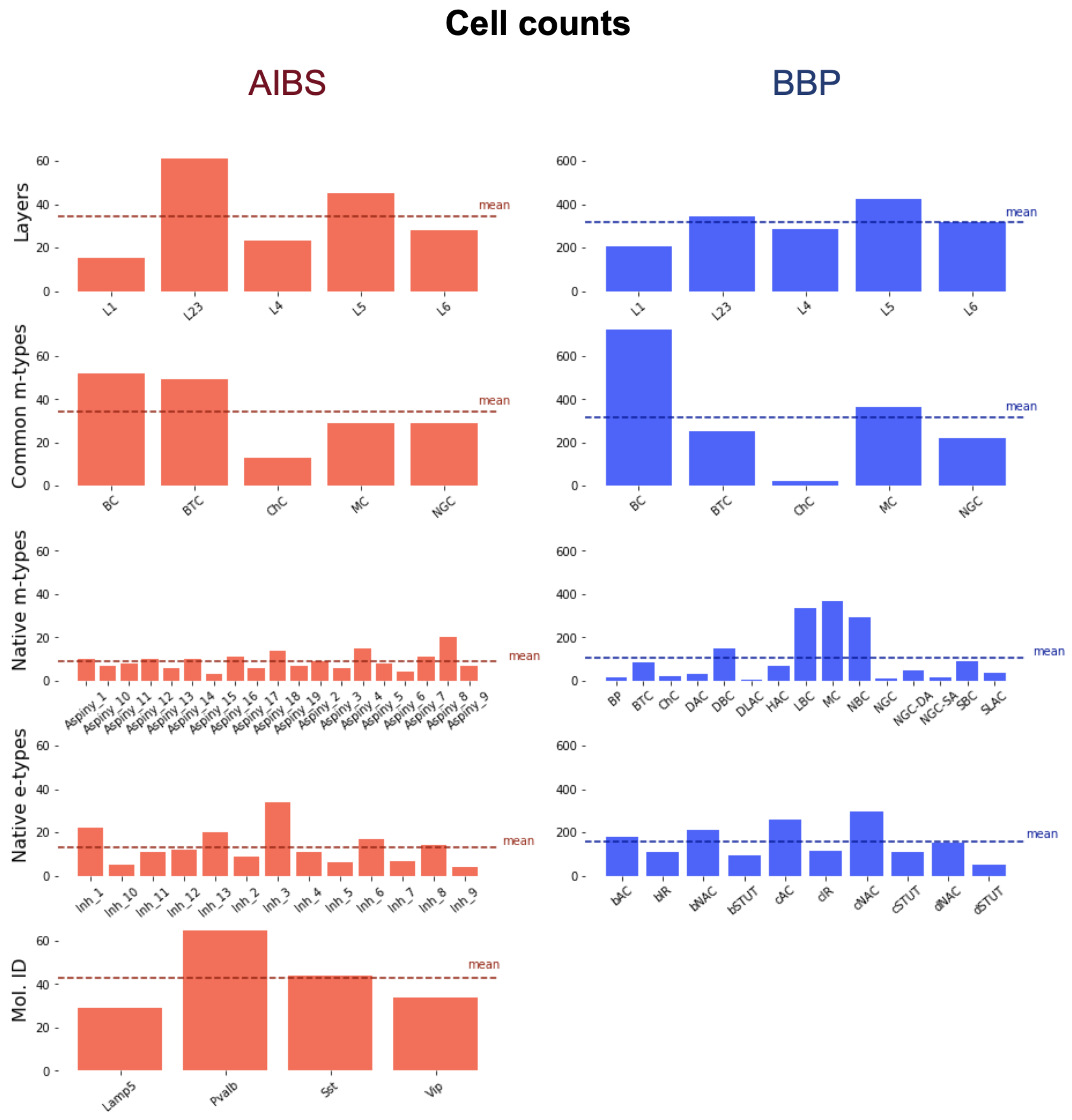


**S5 Figure:** Dataset cell composition according to different labelling systems. Cell counts were grouped by Cortical layer, common m-type (as defined in methods), native m-types, native e-types and molecular ID (when available). AIBS on the left and BBP on the right. Dashed lines represent the mean value for each case.

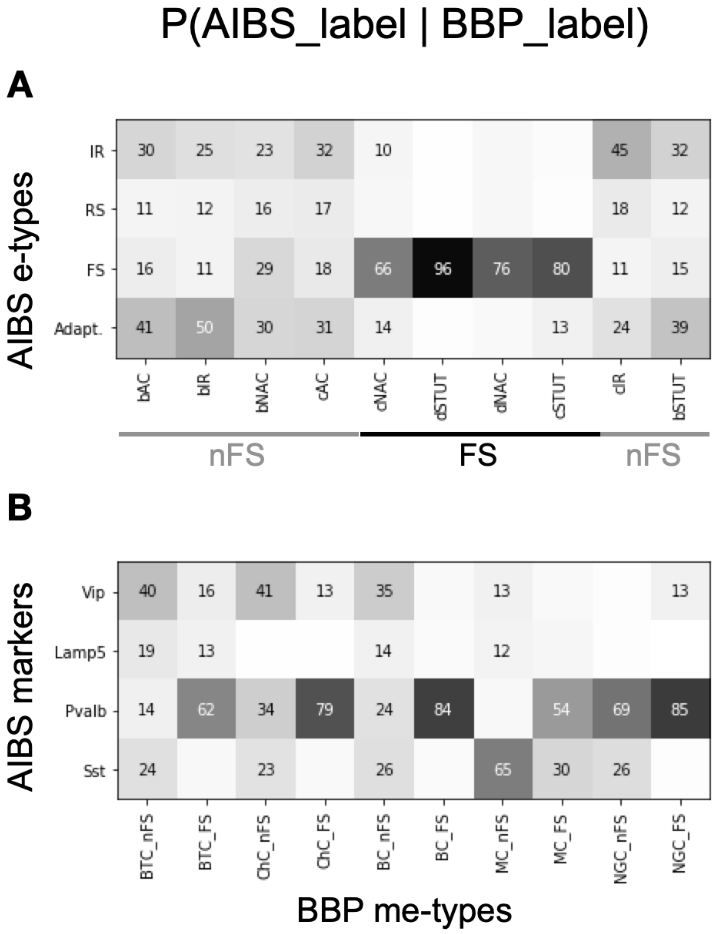


**S6 Figure:** E-types mapping and effect on molecular ID. **A.** Mapping results when the pipeline was applied using the e-types general description provided by Gouwens et al., 2019 for alpha optimization. AIBS neurons were assigned to one of the four following "common" e-types: Irregular spiking (IR), regular spiking (RS), fast spiking (FS) and adapting (Adapt.). Mapping was done between these "common" e-types and BBP e-types. **B.** Mapping results between molecular ID from AIBS dataset and BBP me-types with e-types labelled as either FS or nFS. Exactly the same morphologies are used to build me-models for both e-types.
